# Simultaneous Profiling of the Blood and Gut T and B Cell Repertoires in Crohn’s Disease and Symptomatic Controls Illustrates Tissue-specific Alterations in the Immune Repertoire of Crohn’s Disease Patients

**DOI:** 10.1101/2024.09.22.614337

**Authors:** Aya K. H. Mahdy, Valentina Schöpfel, Gert Huppertz-Hauss, Gøri Perminow, Florian Tran, Corinna Bang, Johannes Roksund Hov, May-Bente Bengtson, Petr Ricanek, Randi Opheim, Tone Bergene Aabrekk, Trond Espen Detlie, Vendel A. Kristensen, Mathilde Poyet, Marte Lie Høivik, Andre Franke, Hesham ElAbd

**Author notes:** The authors wish it to be known that, in their opinion, the first two authors should be regarded as joint first authors. Shared senior authors (Andre Franke, Hesham ElAbd).

## Abstract

<u>C</u>rohn’s <u>d</u>isease (CD) is a clinical subset of inflammatory bowel disease that is characterized by patchy transmural inflammation across the gastrointestinal tract. Although the exact etiology remains unknown, recent findings suggest that it is a complex multifactorial disease with contributions from the host genetics and environmental factors such as the microbiome. We have shown that the T cell repertoire of CD patients harbors a group of highly expanded T cells which hints toward an antigen-mediated pathology. To profile the immunological signature of CD at a high-resolution we simultaneously profiled the αβ and γδ T cell repertoire in addition to the B cell repertoire of both the blood and the colonic mucosa for 27 treatment-naïve CD patients and 27 age-matched symptomatic controls. Regardless of disease, we observed multiple physiological differences between the immune repertoire of blood and colonic mucosa. Additionally, by comparing the repertoire of CD patients relative to controls, we observed different alterations that were only detected in the blood or colonic mucosa. These include a depletion of mucosal-associated invariant T (MAIT) cells in the blood repertoire, an expansion of *TRAV29/DV5-TRAJ5*^+^ clonotypes and a significant depletion of multiple *IGHV3-33-IGHJ4^+^*and *IGHV3-33-IGHJ6^+^* clonotypes in the blood and gut IGH repertoire of CD patients. In conclusion, our findings highlight the importance of studying the immune repertoire in a tissue-specific manner and the need to profile the T and B cell immune repertoire of gut tissues.

## Introduction

Inflammatory <u>b</u>owel <u>d</u>isease (IBD) is a chronic, remitting inflammatory condition of the gut that is clinically subclassified into two main entities, <u>C</u>rohn’s <u>d</u>isease (CD) and <u>u</u>lcerative <u>c</u>olitis (UC). CD is characterized by severe, transmural inflammation that is observed in patches across the entire gastrointestinal tract. In contrast, UC is observed in the colon and rectum with a continuous superficial inflammation that is restricted to the mucosa and the submucosal layer. Dysregulated immune responses have been shown to be a central theme in CD, for example, we have recently shown that yeast specific CD4^+^ T cells exhibit a more cytotoxic phenotype in CD^1^. Furthermore, we have also identified a group of type II like natural killer T (type II NKT) cells that we referred to as <u>C</u>rohn’s <u>a</u>ssociated invariant <u>T</u> (CAIT) cells which are more expanded in CD patients relative to healthy controls and UC patients^2^.

A key determinant of the antigenic specificity of a T cell is its <u>T</u> <u>c</u>ell receptor (TCR), which is generated by a somatic recombination process known as V(D)J in which multiple gene segments encoding the different chains of a TCR recombine to generate an enormous diversity of TCRs. In humans, four TCR chains are utilized to form TCRs, namely, α, β, γ, and δ TCR chains. Generally, α and β chains dimerize to form TCRs observed in αβ T cells, and γ and δ chains dimerize to form TCRs observed in γδ T cells. Based on the class of antigens they recognize; T cells can be classified into conventional T cells, which recognize peptides presented by the <u>h</u>uman leukocyte <u>a</u>ntigen (HLA) proteins, and unconventional T cells which recognize a wider set of non-peptide antigens. Conventional T cells are αβ T cells that recognize peptides presented by HLA-I and HLA-II proteins. Despite this, there is a wide spectrum of unconventional αβ T cells such as <u>m</u>ucosal-<u>a</u>ssociated invariant <u>T</u> cells (MAIT) which recognize vitamin B metabolites presented by <u>M</u>HC-related protein 1 (MR1)^3^. Conversely, γδ T cells are generally unconventional, recognizing a versatile set of non-peptide antigens such as phosphoantigens presented by butyrophilins^4^, sulphatide presented by CD1d^5^, and different stress markers^6^.

Similar to T cells, B cell receptors (BCR) are the main determinant of the antigenic specificity of B cells. BCRs are also generated by V(D)J recombination which generates three main antigen-binding chains, namely, Ig heavy chain (IGH) and two Ig light chains, κ (IGK) and λ (IGL). In addition, the heavy chain can have one of five main constant regions: μ, γ, α, δ, ε which are used in Immunoglobulin M (IgM), G (IgG), A (IgA), D (IgD), and E (IgE) antibodies, respectively. Although BCRs and TCRs are generated by a similar process, *i.e.* V(D)J recombination, and have considerable similarities, BCRs differ in three main features. First, BCR recognizes their antigens directly without any need for antigen presentation by a presenting molecule such as HLA, CD1d, or MR1. While some γδ T cell subsets can recognize their antigens directly without a presenting molecule^7^, most T cells are restricted by their presenting molecule. Second, due to mechanisms such as somatic hypermutation, the genetic sequence of BCRs keeps changing after the formation of functional BCR and antigen stimulation. On the other hand, once T cells egress from the thymus their TCRs are fixed and will not change. Lastly, BCR produces antibodies that have a much higher affinity to their antigenic target in comparison to TCRs.

The repertoire of BCRs, which is a collection of unique BCRs in a population or a collection of B cells, can be profiled by sequencing the V(D)J recombination product encoding for these receptors, through B cell receptor repertoire sequencing (BCR-Seq). Using this framework, Bashford-Rogers and colleagues^8^, have profiled the B cell repertoire of six different autoimmune diseases showing an increase in clonality of IgA in CD and systemic lupus erythematosus, additionally, the authors showed a skewing in the frequency of different IGHV genes in these different diseases. However, these results were obtained using blood samples which might not be able to recapitulate the local immune repertoire of the gut. Indeed, the correlation between the B cell repertoire in the gut and blood remains understudied, for example, Weisel and colleagues^9^ have recently identified a new distinct subset of gut specific B cells. These results highlight the importance of profiling the local repertoire of affected tissues. In this study, we profiled the αβ and γδ T cell repertoires, as well as the B cell repertoire of paired blood and gut samples (obtained using biopsies) of 27 CD patients and age-matched symptomatic controls (total =108 samples).

## Results

### Cohort description

We used samples derived from the inception-cohort inflammatory <u>b</u>owel disease in <u>S</u>outh- <u>E</u>astern <u>N</u>orway III (IBSEN-III)^10^. Specifically, we used DNA extracted from EDTA blood tubes and biopsies derived from colonic mucosa (gut biopsies hereafter) collected at the time of diagnostic endoscopic procedure to profile the full T and B cell repertoire (**Material and Methods**) of 27 active (inception; recently diagnosed) treatment-naïve CD patients and 27 age-matched symptomatic controls (*i.e.* patients referred with suspicion of IBD but whose diagnostic work-up, including endoscopy and histology, was normal) (**Table 1**). For the 27 controls, biopsies were derived from non-inflamed tissues from the left-side of the colon, while for CD patients, we utilized 11 biopsies from inflamed and 16 from non-inflamed colons (left-side). Following the immune profiling we aimed at investigating two questions, first, the physiological differences in the immune repertoire of these two tissues, second, similarities and differences in CD induced immune changes that are either detected at the local gut immune repertoire and/or at the blood repertoire level.

**Table 1:** Phenotypic and clinical parameters of CD and symptomatic controls where the T and B cell repertoire of blood and gut tissue was profiled. ^†^ The two patients with Upper GI tract disease location also had disease observed at the ileocolon, *i.e.* they were L3+L4 cases. ^#^ Represents cases with <u>n</u>on- <u>s</u>tricturing, <u>n</u>on-<u>p</u>enetrating (NSNP) disease.

|  | *Crohn's disease* (n=27) | *Symptomatic controls* (n=27) |
| --- | --- | --- |
| *Age* | 27.07 ± 6.13 (18-39) | 27.81 ± 6.28 (19-38) |
| *Female percentage* | 44% (n=12) | 59% (n=16) |
| *Disease location* | L1 (Ileum): 6 probands<br>L2 (Colon): 7 probands<br>L3 (Ileocolon): 14 probands<br>L4 (Upper GI tract): 2<sup>†</sup> probands | N/A |
| *Behavior overtime* | B1 (NSNP<sup>#</sup>): 16 probands<br>B2 (Stricturing): 7 probands<br>B3 (Penetrating): 2 probands | N/A |

### Different immune receptor loci exhibit a different degree of overlap between the blood and gut, and show different degrees of interindividual overlap

Across all samples and profiled immune receptor chain, namely, TRA, TRB, TRG, TRD, IGH, IGK, and IGL, 2,034,407 unique productive clonotypes, *i.e.* unique V(D)J arrangement that did not contain a frameshift or a stop codon, were identified. Among them, 852,491 were observed only in blood samples (∼42%) while 1,117,271 (∼55%) were identified from gut tissues and only 64,645 (∼3%) clonotypes were detected in the repertoire of both compartments (**Fig. 1A**). For two donors, (1 CD and 1 control) we observed a very small number of clonotypes which also correlated with a low-quality input DNA, subsequently, samples derived from these two patients were excluded from all analyses. From an immune receptor chain perspective, there was a clear separation between the immune repertoire of gut tissues and blood with a more diverse T cell repertoire in blood and a more diverse B cell repertoire in the gut (**Fig. 1B**). Specifically, the TRA and the TRG showed higher diversity in the blood compared to gut tissues, while the IGH showed substantially greater diversity in the gut than in the blood (**Fig. 1B**). To explore this further, we calculated the clonal expansion index of each locus in both blood and gut samples (**Fig. 1C**). We did not observe any significant difference in the clonal expansion index of the T cell repertoires of gut and blood, across the four loci (**Fig. 1C**). However, there was a significant difference in the clonal expansion of the B cell repertoire (IGH, IGK, IGL), with the blood repertoire showing a higher degree of clonal expansion compared to the local gut B cell repertoire (**Fig. 1C**).

**Figure 1:**
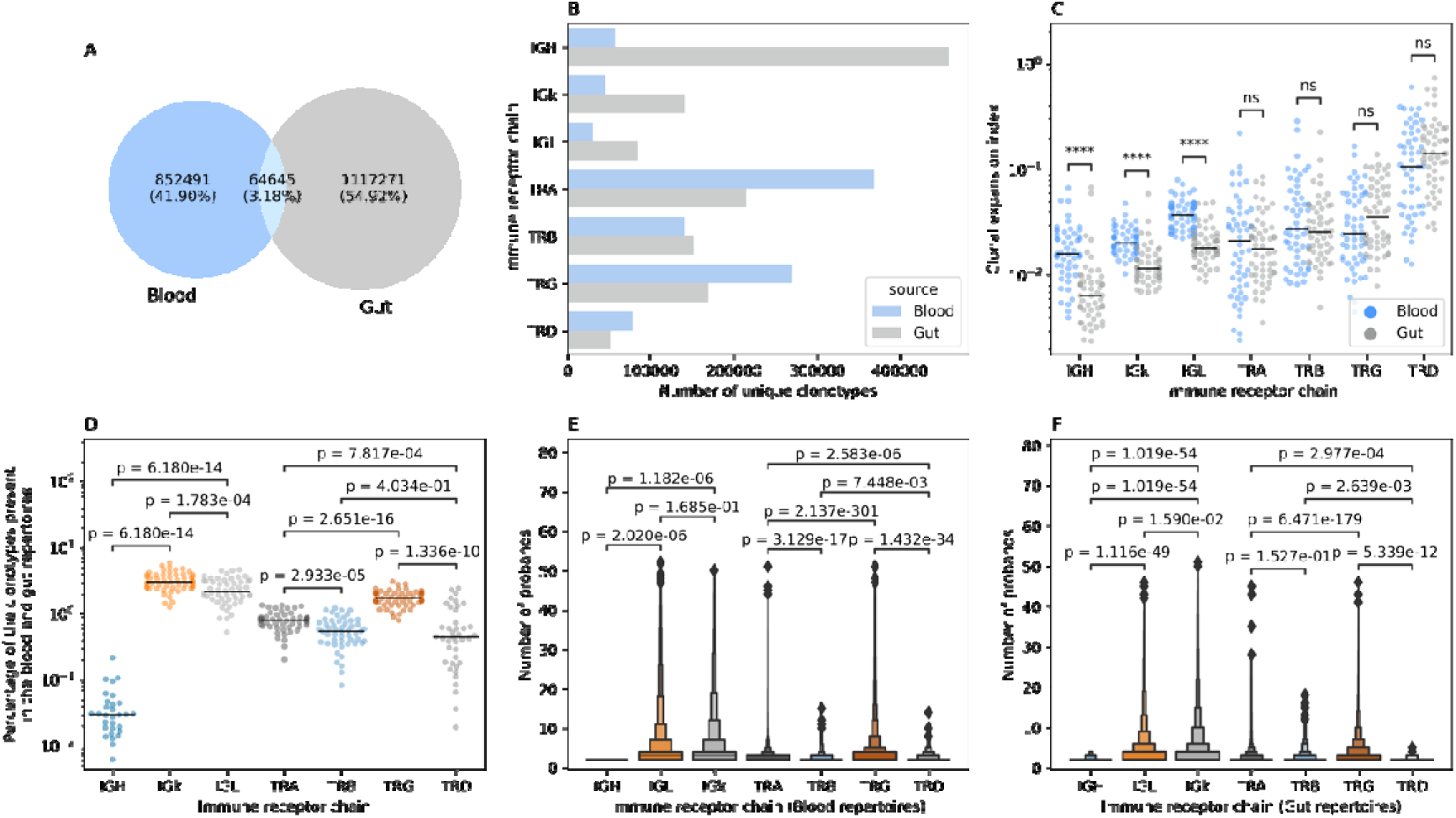
Overlap between the blood and gut repertoire among the seven different immune receptor loci. **(A)** Overlap in productive clonotypes between the blood and gut repertoire among all immune receptor loci. **(B)** The number of clonotypes observed by each immune receptor gene in the gut and the blood. **(C)** Clonal expansion index of clonotypes observed in the blood and the gut repertoire of different immune receptor genes. **(D)** Percentage of overlapped clonotypes between clonotypes detected in the blood and gut repertoire over the union of clonotypes observed in the gut and the blood repertoires of the different loci. **(E)** and **(F)** boxen plot representation of the degree of publicity, *i.e.* the number of probands having the same clonotype in their repertoire, of public blood and public gut clonotypes, respectively. In **(C)**, **(D)**, **(E),** and **(F)**, statistical comparisons were conducted using the two-sided Mann-Whitney-Wilcoxon test. In this figure and all subsequent figures unless stated otherwise, the asterisks are used to represent the P-values, with n.s. donating non-significant (P>0.05), * represents P 0.05, ** donates, P 0.01, *** means P 0.001 and lastly, **** designates a P 0.0001.

We next quantified the overlap between the blood and colon repertoires at the intra-individual level as well as the overlap between the blood and gut repertoires across different individuals (inter-individual level variation). Among the B cell repertoire chains, IGK showed the highest degree of intra-individual overlap between the blood and the gut immune repertoires (median ∼3%). In contrast, IGH showed the lowest overlap between these two compartments (median ∼0.03%) (**Fig. 1D**). IGL showed an intermediate level of overlap between the blood and gut (median ∼2.15%) which is significantly lower than IGK but nearly 71 times higher than IGH (**Fig. 1D**). A similar pattern was observed for the T cell repertoire chains. The TRG showed the highest degree of overlap (median ∼1.68%), followed by the TRA loci (median ∼0.78%) (**Fig. 1D**). The least overlap was observed for the TRB and TRD loci, with a median of ∼0.53% and ∼0.43%, respectively (**Fig. 1D**).

A similar pattern was observed at the inter-individual level for the blood repertoire, where the light chains of the BCR, *i.e.* IGL, and IGK, are shared among a higher number of donors, with a median of 3 probands, relative to the heavy chain, *i.e.* IGH, with a median of 2 (**Fig. 1E**). It is worth noticing that this analysis focused on public clonotypes which, by definition, are clonotypes observed in two or more probands. At the T cell repertoire level, TRG was the most public among the four TCR chains with a mean of ∼3.5 probands, followed by TRA (mean ∼2.56 proband), then TRD (∼2.35 proband) and lastly TRB (mean ∼2.29 probands) (**Fig. 1E**). Across the four chains the median was 2, suggesting that these differences are driven by sub-populations of highly public clonotypes, that are present in many individuals. These populations were more frequent in the TRG repertoire followed by the TRA, the TRD and lastly, the TRB repertoire. A similar pattern was observed at the gut immune repertoire with the light chains being more public relative to the heavy chain, however, IGK clonotypes were slightly more public than IGL clonotypes with a mean of 3.97 and 3.74, respectively (**Fig. 1F**). At the T cell repertoire level, there was not any significant difference in the degree of publicity between the TRA and TRB repertoires, with the TRG repertoire exhibiting the highest degree of publicity relative to all other TCR chains (**Fig. 1F**).

### The blood and gut immune repertoire significantly differ in the utilized V and J genes

Next, we analyzed the V gene usage in the immune repertoire of blood and gut tissues. Across the TRA repertoire we observed a significant difference in gene usage, with *TRAV1- 2*, *TRAV26-2*, *TRAV27-2* and *TRAV39* showing significantly higher frequencies in blood relative to gut tissues (**Fig. 2A**). On the contrary, *TRAV8-1*, *TRAV8-2*, *TRAV8-3*, *TRAV8-4*, and *TRAV8-6* showed higher frequency in gut tissues compared to blood (**Fig. 2A**). This prominent difference was not observed at the TRB repertoire with only *TRBV18* having higher frequency in gut repertoire and *TRBV4-1*, *TRBV6-05*, *TRBV27*, *TRBV7-9* and *TRBV7-2* showing higher frequency in the blood repertoire (**Fig. 2B**). Some of these V-genes are known to be preferentially used by different subsets of unconventional T cells, for example, *TRAV1-2* which encodes the α7.2 chain, is commonly utilized by MAIT cells. Similarly, *TRAV26-2 and TRBV7-9*^11^ as well as *TRBV4-01*^12^ have been observed in CD1c restricted T cells. These results suggest that within the α/β T cell population, different cellular subsets preferentially reside in these two anatomical compartments. The same findings were observed in the γ/δ T cell repertoire, with the frequency of *TRGV10* being higher in the gut repertoire and that of *TRGV9* being higher in the blood repertoire (**Fig. 2C**). Within the TRD repertoire, *TRDV1* and *TRDV3* had a higher frequency in gut tissues relative to blood while *TRDV2* dominated the blood repertoire (**Fig. 2D**). TRGV9 preferentially pairs with the *TRDV2* to form the γ9/δ2 subset which is the main subset of γ/δ T cells observed in blood. TRDV1^+^ clonotypes, *i.e.* δ1^+^ T cells, are the main subset of γ/δ T cells that are commonly observed in gut tissues.

**Figure 2:**
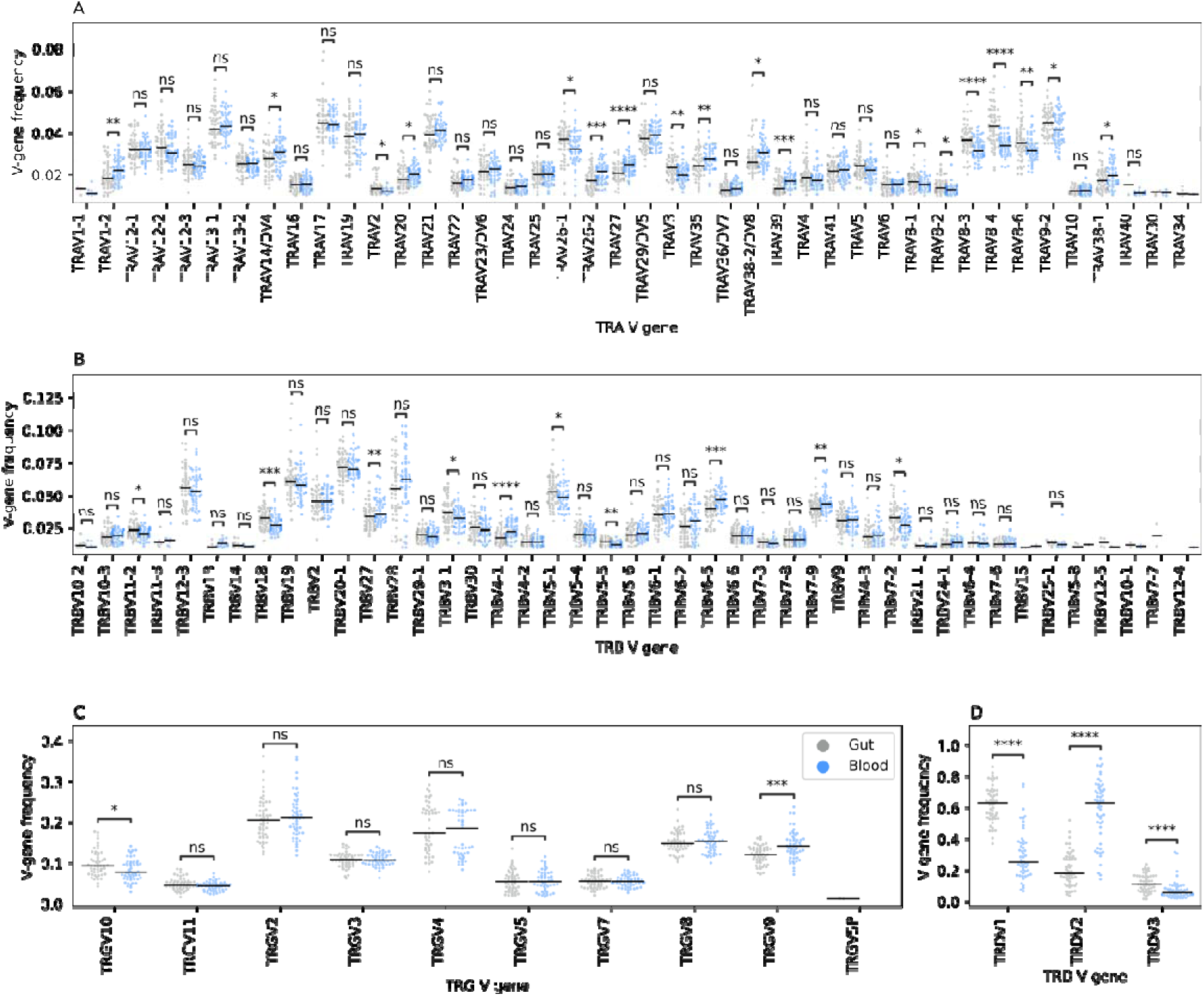
the relative frequency of different V genes among the four TCR chains in the gut and blood immune repertoires. (**A**) depicts the differences at the TCR alpha (TRA) chain, while (**B**) depicts the differences at the TCR beta (TRB) chain, (**C**) at the TCR gamma (TRG) chain and lastly, (**D**) at the TCR delta (TRD) chain. Across all panels, blackline represents the median, we also used the two-sided Mann-Whitney-Wilcoxon test to compare the frequency of each V-gene segment among the two anatomical compartments. Additionally, we filtered V-genes with a frequency less than 0.01, *i.e.* 1%, from the analysis as well as V-gene segments with more than 1% frequency in less than 10 samples from any statistical comparisons.

Across the three B cell receptor chains, namely, IGH, IGK and IGL, we observed significant differences in V-gene usage between the blood and gut repertoires (**Fig. S1**). Specifically, within the IGH repertoire, where *IGHV3-74*, *IGHV3-7* and *IGHV3-23* were found at significantly higher frequencies in gut tissue compared to blood, while *IGHV1-18, IGHV1-69, IGHV3-21, IGHV3-34 and IGHV3-30-3* were more frequent in blood than in gut tissues (**Fig. S1A**). Given that we analyzed the joint repertoires of CD patients and controls, these findings suggest a physiological difference between blood and gut repertoires. Suggesting that CD might lead to tissue-specific changes in the immune repertoire, or that disease-induced changes could vary in magnitude between different tissues.

### Crohn’s disease induced multiple changes in the blood TCR repertoire that are not detected in the gut repertoire

To identify CD induced changes in the blood TCR repertoire, we first compared the frequency of V-genes between CD patients and controls (**Fig. 3A**). We observed a significant difference in the frequency of three V-genes: *TRAV23/DV6*, *TRAV4*, and *TRAV6* (**Fig. 3A**). We then investigated the expansion of clonotypes harboring these V-gene segments in both CD patients and controls. Although trends were detected, with a depletion of *TRAV23/DV6*^+^ clonotypes (**Fig. 3B**), and an expansion of *TRAV4*^+^ (**Fig. 3C**) and TRAV6^+^ clonotypes in CD (**Fig. 3D**), none of these trends was significant, likely due to the small sample size.

**Figure 3:**
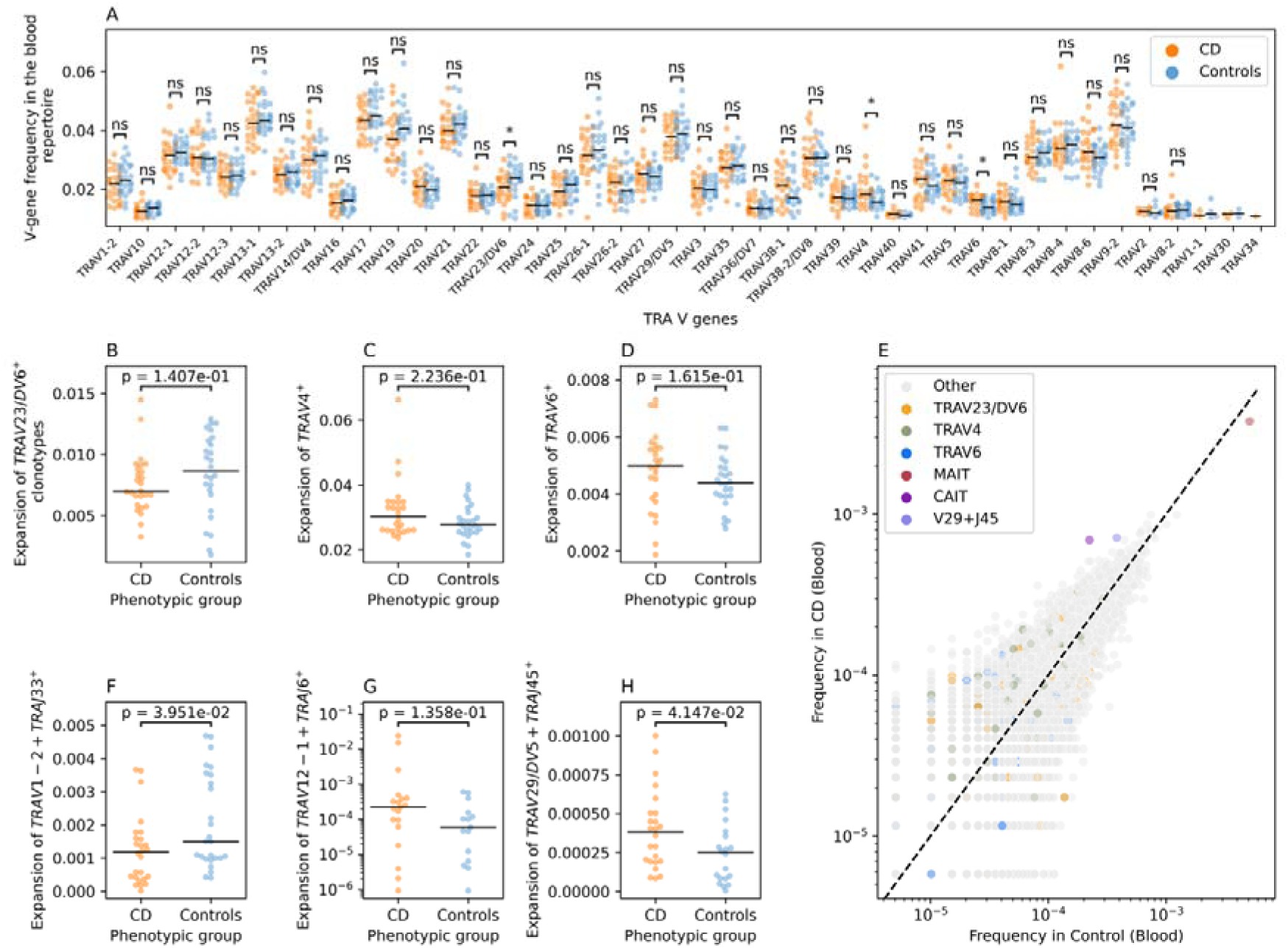
CD-induced changes on the blood TRA repertoire of CD patients and controls. (**A**) a comparison between the frequency of different TRA V-genes in CD patients and controls. (**B**) shows the difference in the expansion of TRAV23/DV6^+^ clonotypes in CD and Controls, while (**C**) illustrates the same relationship but for TRAV4^+^ clonotypes and (**D**) for TRAV6^+^. (**E**) shows the frequency of different VJ-length groups in the TRA blood repertoire of controls and CD patients. (**F**) shows the expansion of the TRAV1-2+TRAJ33+L12 which is used as a proxy for MAIT cells in CD patients and Controls, while (**G**) depicts the same relationship but for TRAV12- 1+TRAJ+L15 group which is a superset of CAIT cells and lastly, (**H**) illustrate the expansion of TRAV29/DV5- TRAJ45+L16 group in the blood of CD patients relative to controls. Across all panels except (**E**), we used the two-sided Mann-Whitney-Wilcoxon test to compare the frequency of each V-gene segment between CD patients and controls.

To obtain a more detailed view, we categorized clonotypes into refined groups by combining unique V and J genes with specific CDR3 amino acid lengths, creating VJ-CDR3 length groups. This approach revealed several alterations, including a depletion of TRAV1- 2+TRAJ33^+^ clonotypes with a CDR3 length of 12 amino acids (TRAV1-2+TRAJ33+L12) in the blood repertoire of CD patients (**Fig. 3E**). Since the TRAV1-2+TRAJ33 rearrangement is commonly observed in MAIT cells, we labeled this arrangement as MAIT cells. We observed a significant depletion of MAIT cells (defined as: TRAV1-2+TRAJ33+L12) in the blood of CD patients, consistent with our previous findings and those of others^2,13^ (**Fig. 3F**).

We also observed an expansion of two other VJ-CDR3 length groups in CD patients compared to controls, namely, TRAV12-1+TRAJ6+L15 and TRAV29/DV5+TRAJ45+L16 (**Fig. 3E**). The TRAV12-1+TRAJ6+L15 group includes a subset of unconventional T cells we previously identified as Crohn’s-associated invariant T cells (CAIT cells)^2^. CAIT cells are characterized by a semi-invariant TCR-alpha chain comprising *TRAV12-1* and *TRAJ6* combination and a CDR3 amino acid sequence represented by the motif CVV**A*GGSYIPTF, where the asterisk represents any amino acids^2^. We used TRAV12- 1+TRAJ6+L15 as a proxy for CAIT cells. Although a trend toward expansion was observed (**Fig. 3G**), it was not statistically significant, likely due to our proxy definition which was not specific enough with regard to the established CAIT cells motif. In the case of the TRAV12- 1+TRAJ6+L15 VJ length group, we know that it contains other subsets beside the CAIT cells, which dampen the CAIT signal. However, for the sake of fair comparison between the VJ-Length groups, we did not used the more refined definition of CAIT cells to compare its expansion in CD patients and controls. Lastly, we observed a significant expansion of TRAV29/DV5+TRAJ45+L16 clonotypes in the blood of CD patients. To the best of our knowledge, the expansion of this group in the blood of CD patients has not been previously reported and could suggest a novel aspect of the immune **response in CD patients.**

Interestingly, the effect observed in the blood TCR repertoire were not detected in the gut TRA repertoire (**Fig. S2**). For example, while comparing the V-gene usage, we observed a reduction in the frequency of *TRAV12-1^+^* clonotypes and an increase in TRAV8-6^+^ clonotypes in the blood of CD patients relative to controls (**Fig. S2A**). Nonetheless, we did not detect significant changes in the expansion or depletion of these clonotypes in the gut repertoire of CD patients and controls (**Fig. S2B-S2C**). Furthermore, candidate clonotype groups identified in the blood repertoire, namely, TRAV1-2+TRAJ33+L12, TRAV12- 1+TRAJ6+L15 and TRAV29/DV5+TRAJ45+L16, did not show a differential degree of expansion in CD patients relative to controls at the gut repertoire **Fig. S2D-2F**, respectively.

Additionally, by comparing the frequency of different VJ-CDR3 length groups, we could not identify a group that was significantly different between CD patients and controls.

Despite the small sample-size, CD-associated changes were identified in the TRA repertoire, but these did not translate to other TCR chains. For example, we could not detect any significant changes at the TRB repertoire neither in blood nor at the gut level (**Fig. S3**). Similarly, we failed to detect any significant differences in the γδ T cell repertoire of blood and gut (**Fig. S4**).

### Crohn’s disease is associated with changes in the IGH repertoire

For BCR repertoire analysis, we examined the IGK and IGL light chains in both blood and gut but found no significant differences between CD patients and controls (**Fig. S5-S8**). Subsequently, we focused on the immunoglobin heavy chain repertoire (IGH). By comparing the IGH V-gene frequency in the blood repertoire of CD patients and controls we observed a significant reduction in the frequency of *IGHV3-33*^+^ in the blood repertoire of CD patients relative to controls (**Fig. S9**). Subsequently, we binned clonotypes into groups using the same VJ-length approach discussed above (**Fig. 4A**). This enabled us to identify multiple groups that had differential frequency in CD patients relative to controls. One of these groups was IGHV3-30+IGHJ6+L24, which had a higher frequency in cases relative to controls, however, by comparing the expansion of this group between CD patients and controls we could not detect any significant difference (**Fig. 4B**). We also observed a significant depletion of a subset of *IGHV3-33+IGHJ4*^+^ and *IGHV3-33+IGHJ6^+^* clonotypes, specifically, IGHV3-33+IGHJ4+L18, IGHV3-33+IGHJ4+L19, IGHV3-33+IGHJ4+L16, IGHV3-33+IGHJ6+L25, and IGHV3-33+IGHJ4+L15 (**Fig. 4A**). These clonotypes were significantly depleted in CD patients relative to controls (**Fig. 4C**). We also observed a group of *IGHV3- 23^+^* clonotypes that had higher frequency in CD and controls and showed different frequency between the two groups (**Fig. 4A**).

**Figure 4:**
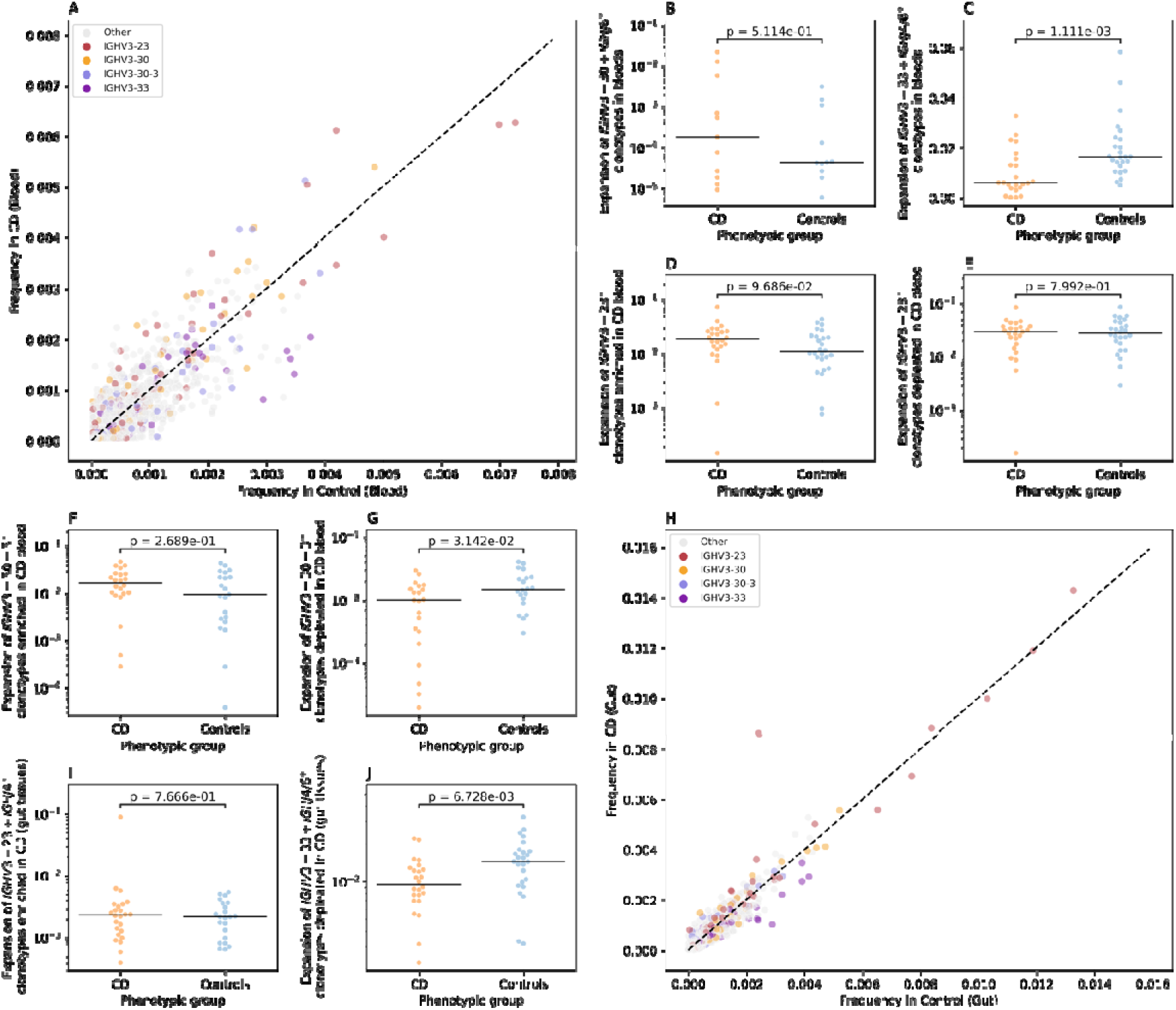
CD-associated changes in the IGH repertoire of colonic mucosa and peripheral blood. (**A**) the differential frequency of VJ-length groups in the blood repertoire of CD patients and controls. (**B**) shows the expansion of IGHV3-30+IGHJ6+L24 clonotypes in the blood repertoire of CD patients and controls, while (**C**) illustrate the depletion of IGHV3-33+IGHJ4+L18, IGHV3-33+IGHJ4+L19, IGHV3-33+IGHJ4+L16, IGHV3-33+IGHJ6+L25, and IGHV3-33+IGHJ4+L15 clonotypes in the blood repertoire of CD patients. (**D**) and (**E**) shows the expansion of IGHV3-23^+^ with a frequency higher than 0.002 in the blood of CD and controls, with (**D**) depicting the expansion for VJ-length groups with a higher frequency in CD and (**E**) showing the expansion of VJ- length group with higher frequency in controls relative to CD. (**F**) and (**G**) depict the expansion of IGHV3-30-3 VJ- length groups with a frequency higher than 0.002 in the blood of CD and controls with (**F**) showing the expansion of IGHV3-30-3^+^ clonotypes with a higher frequency in CD relative to controls and (**G**) depicting the difference in the expansion of VJ length groups with higher frequency in controls. (**H**) the differential frequency of VJ-length groups in the Gut repertoire of CD patients and controls. (**I**) the expansion of IGHV3-23+IGHJ4+L22 clonotypes in the gut IGH repertoire of CD and controls. Lastly (**J**), depicts the depletion of IGHV3-33+IGHJ4+L18, IGHV3- 33+IGHJ4+L19, IGHV3-33+IGHJ4+L16, IGHV3-33+IGHJ6+L25, and IGHV3-33+IGHJ4+L15 clonotypes in the gut IGH repertoire of CD patients.

To disentangle this further, we selected *IGHV3-23^+^* VJ-length group with a frequency above 0.002 (*i.e.* 0.2%) in cases and controls. After that we compared the expansion of subgroups with higher frequency in CD patients (**Fig. 4D**) or in controls (**Fig. 3E**), which showed no significant difference in neither of these groups. The same framework was used to analyze the IGHV3-30-3^+^ clonotypes, where VJ-length group with higher frequency in CD relative to controls did not show a significant difference in the expansion (**Fig. 3F**). On the other side, IGHV3-30-3^+^ VJ length groups with a higher frequency in control, illustrated a significant depletion in the blood repertoire of CD patients relative to controls (**Fig. 3G**)

Subsequently, we repeated the VJ length group analysis using the gut IGH repertoire (**Fig. 4H**). We observed a higher frequency from a specific VJ-length group, namely, IGHV3- 23+IGHJ4+L22 in CD patients relative to controls (**Fig. 4H**). Nonetheless, by comparing the expansion of this group of clonotypes in cases relative to controls, we did not observe any significant differences with only one CD samples showing extremely high level of expansion (**Fig. 4I**). Lastly, we aimed at investigating the IGHV3-33 depletion signal observed at the blood repertoire (**Fig. 4C**) in the gut repertoire. To this end, we compared the expansion of *IGHV3-33+IGHJ4*^+^ and *IGHV3-33+IGHJ6^+^* clonotypes VJ-length groups defined above in the gut IGH repertoire of CD patients and controls which confirmed the depletion of these clonotype groups at the local gut repertoire.

## Discussion

To study Crohn’s disease effects on different immune repertoire compartments, we profiled the T and B cell immune repertoires in blood and colonic mucosa from 27 CD patients and 27 age-matched symptomatic controls from the IBSEN III cohort^10^. This enabled us to identify physiological differences between the blood and gut repertoire irrespective of the disease status. In addition to identifying disease-alterations that were observed either at the gut and/or the blood. Nonetheless, before discussing the immunological and clinical implications of our findings, we would like to discuss methodological and technical aspect influencing our findings. The quality and quantity of the input DNA were a fundamental challenge for profiling the immune repertoire from mucosal biopsies where the utilization of low-quality DNA usually fail to produce trustworthy immune profiles. For all the analyses reported here, we removed the immune repertoire of two samples because of their initial low quality input DNA.

Other aspects beyond DNA quality needs to be taken into consideration when profiling the immune repertoire across tissues, namely, cellular composition and spatial differences. Peripheral blood contains a large fraction of T (70%) and B (15%) cells^14^, however, theses proportions vary across tissues. Although we used an equal amount of DNA (120 ng) for profiling the blood and the gut repertoires, differences in cellular composition between these two tissues, does not equate to equal numbers of T and B cells in the starting material. Additionally, spatial heterogeneity within tissues can impact the results of the immune profiling. To minimize this variability, we focused on biopsies from the left-side of the colon, including both inflamed and non-inflamed biopsies from CD patients and controls. However, this approach may have introduced heterogeneity in the collected gut biopsies limiting our ability to detect CD-associated changes in the gut repertoire.

Despites these technical aspects, our analysis revealed that changes were primarily observed in the TRA repertoire. While other studies have reported alterations in the TRB^15^ and the TRG^16^ repertoires, they involved larger sample sizes, suggesting that analyzing the TRA repertoire might be a more sample-efficient method to identifying disease-associated clonotypes. A key finding in our study was the depletion of MAIT cells (defined using its semi-invariant alpha-chain) in the blood of IBD patients, consistent with previous reports^2,13,17–19^. Using paired measurements of blood and gut samples from the same individual we could show that this effect is only observed in the blood repertoire and not in the gut repertoire. This indicates that this depletion is not due to the migration of MAIT cells to the gut, as CD patients and controls had comparable levels of MAIT cells in their gut tissues.

Beyond the decrease in their abundance, MAIT cells in IBD show a different functional and phenotypic profile that is characterized by a reduction in IFN-γ upon stimulation, in addition to a higher expression of the apoptotic marker annexin V and the exhaustion marker PD-1^19^. The higher expression of proapoptotic markers which was also observed by Hiejima and colleagues^13^, coupled with the expression of exhaustion markers indicate that in IBD, MAIT cells might be decreased because of an exhaustion-induced apoptosis which is observed with chronic T cell exhaustion^20,21^. A recent study by Yasutomi and colleagues^22^ highlighted the pathogenic role of MAIT cells in IBD, showing that MR1 deficient mice have a less severe outcome in an Oxazolone-induced colitis model. Interestingly, the authors also reported that the administration of the MR1 antagonist, isobutyl 6-formyl pterin reduced the colitis severity in the same model as well as decreased the production of cytokines by MAIT cells^22^.

In addition to our work on T cells, we performed an in-depth investigation of the blood and gut B cell repertoire in CD patients and symptomatic controls. This study represents, to the best of our knowledge, the first bulk gut B cell repertoire profiling in CD as well as paired profiling of the blood and gut B cell repertoire. Our findings highlight the importance of studying tissue-specific B cell repertoires, given the low level of overlap observed between the blood and gut B cell repertoire. Although we identified multiple abnormalities in both compartments, the functional implications, *e.g.* the target antigen, remain unknown. To address this, integrating BCR-Seq methods with large-scale antigen screening methods such as <u>ph</u>age-immuno<u>p</u>recipitation sequencing (PhIP-Seq)^23^ could associate antigenic binding to specific B cell expansions and rearrangement.

Our results demonstrate the feasibility and potential of simultaneous T and B cell repertoire profiling using DNA from blood and tissues samples. However, the small sample size included in the study limits its statistical power and our ability to detect disease-associated changes with weaker effects. In addition, our DNA-based approach did not allow for the identification of the B cell receptor isotypes, *e.g.* μ, γ, or α, limiting insights into the biological significance of the identified clonotypes. Lastly, our profiling method did not contain unique molecular identifiers, which hinders our ability to quantify clonal expansion as well as to remove technical artifacts introduced by PCR amplifications. Future studies should include larger sample sizes, incorporate unique molecular identifiers in the amplicon design, and utilize RNA, whenever possible, to capture the V(D)J recombination along with the heavy chain isotype for a more comprehensive profiling.

## Author contributions

A.M., M.L.H, M.P., H.E. and A.F. conceived and outlined the study. A.M. and V.S. conducted the experimental work and prepared the sequencing libraries. H.E. and A.M. analyzed the generated datasets. H.E. and A.M. wrote the manuscript with input from all co-authors. C.O, G.H., G.P., F.T., C.B., J.H., M.B., P.R., R.O., S.A., T.A., T.D., V.K. S.M. and M.H. collected the samples utilized in the current study and provided clinical information about the probands included in the study. All authors reviewed, edited, and approved the final manuscript.

## Ethical approval

The IBSEN III study was approved by the South-Eastern Regional Committee for Medical and Health Research Ethics (Ref 2015/946-3) and performed in accordance with the Declaration of Helsinki. All patients signed an informed consent form prior inclusion in this study and the data were stored in services for sensitive data (TSD) at the University of Oslo.

## Availability of Data

Data are not deposited in a public repository due to data privacy regulations in Norway and our institution. However, data are available upon request, if the aims of the planned analyses are covered by the written informed consent signed by the participants, pending an amendment to the ethical approvals and a material & data transfer agreement between the institutions.

## Material and Methods

### Sample collection

Blood and colon biopsies were collected during the diagnostic work-up of patients included in the IBSEN III study. Patients referred to the hospitals under the suspicion of IBD were invited. Patients who fulfilled the Lennard Jones criteria^24^ for CD were included as CD patients while patients referred with suspicion of IBD but where diagnostic work-up, including endoscopy and histology, was completely normal were included as symptomatic controls. Two mucosal biopsies were collected from the left colon using jumbo biopsy forceps and immersed in Allprotect © (Qiagen) before incubation at 2-8C overnight and then freezing at - 80°C. Blood was drawn in 6 ml EDTA tubes and frozen at -80°C.

### DNA extraction

The blood was collected into EDTA tubes and DNA extraction was performed using *Chemagic STAR* (*Hamilton*) with the *chemagic*™ *DNA Blood 10k Kit H12* from *Revvity* for isolating very high-quality DNA. DNA was isolated from biopsies using a *QIAcube Connect* device and the *DNeasy Blood&Tissue Kit* (Qiagen). Quality control (QC) for the DNA was performed by Genomic DNA (gDNA) ScreenTape Analysis using Agilent TapeStation Systems. The quality of extracted gDNA from EDTA tubes and biopsies ranged from 6.5-9.7 (DIN score). DNA concentration measurements were performed by Invitrogen™ Qubit™ dsDNA-BR (Broad Range) assay kit with qubit fluorometer such as Victor Nivo multimode microplate reader, (PerkinElmer).

### Library preparation, Sample pooling, sequencing, and clonotype identification

The 7genes DNA multiplexing kits from MiLaboratories were used for profiling the full repertoire of T and B cells in gut and blood samples (108 samples prepared from 56 different probands). 120ng of DNA were used for library preparation according to the manufacturer’s instructions. The generated libraries were pooled and sequenced together based on the sequencing recommendation protocols from MiLaboratories kits that were used for library preparation. Libraries prepared using the 7genes Kit from MiLaboratories were prepared as described above, subsequently, these libraries were pooled together and cleaned with magnetic beads (AMPure, ^®^XP Beads protocol, Beckman Coulter, with 1:0.8 sample: beads ratio). Lastly, samples were diluted to a final concentration of ∼2nM and 60 µl of this pool were then submitted for sequencing at the <u>C</u>ompetence <u>C</u>entre for <u>G</u>enomic <u>A</u>nalysis (CCGA) Kiel. Approximately 10% of PhiX was added to this pool which contained 118 samples (108 samples from 56 probands, 8 negative controls, and 2 positive controls) prior to sequencing on a single S4 lane on a NovaSeq 6000 machine using a paired 150bp x2 approach. From these 118 samples, 2,742,957,859 reads were generated with a median of 21,215,996.5 and a mean of 23,245,405.58 reads per samples. Across all samples, a median of ∼ 45.65% and a mean of ∼48% of reads could not be aligned by MIXCR^25^ using the “milab-human-dna-xcr-7genes-multiplex” preset, consequently these reads were not used for clonotype identification. Furthermore, the aligned reads did not have a uniform distribution across all loci with a median of ∼10.9% of reads aligning to the TRA, ∼3.43% to the TRB, ∼52.2% to the TRG, ∼11.55% to the TRD, ∼4.93% to the IGH, ∼5.37% to the IGK and ∼2.28% to the IGL loci.

---

**Figure S1:**
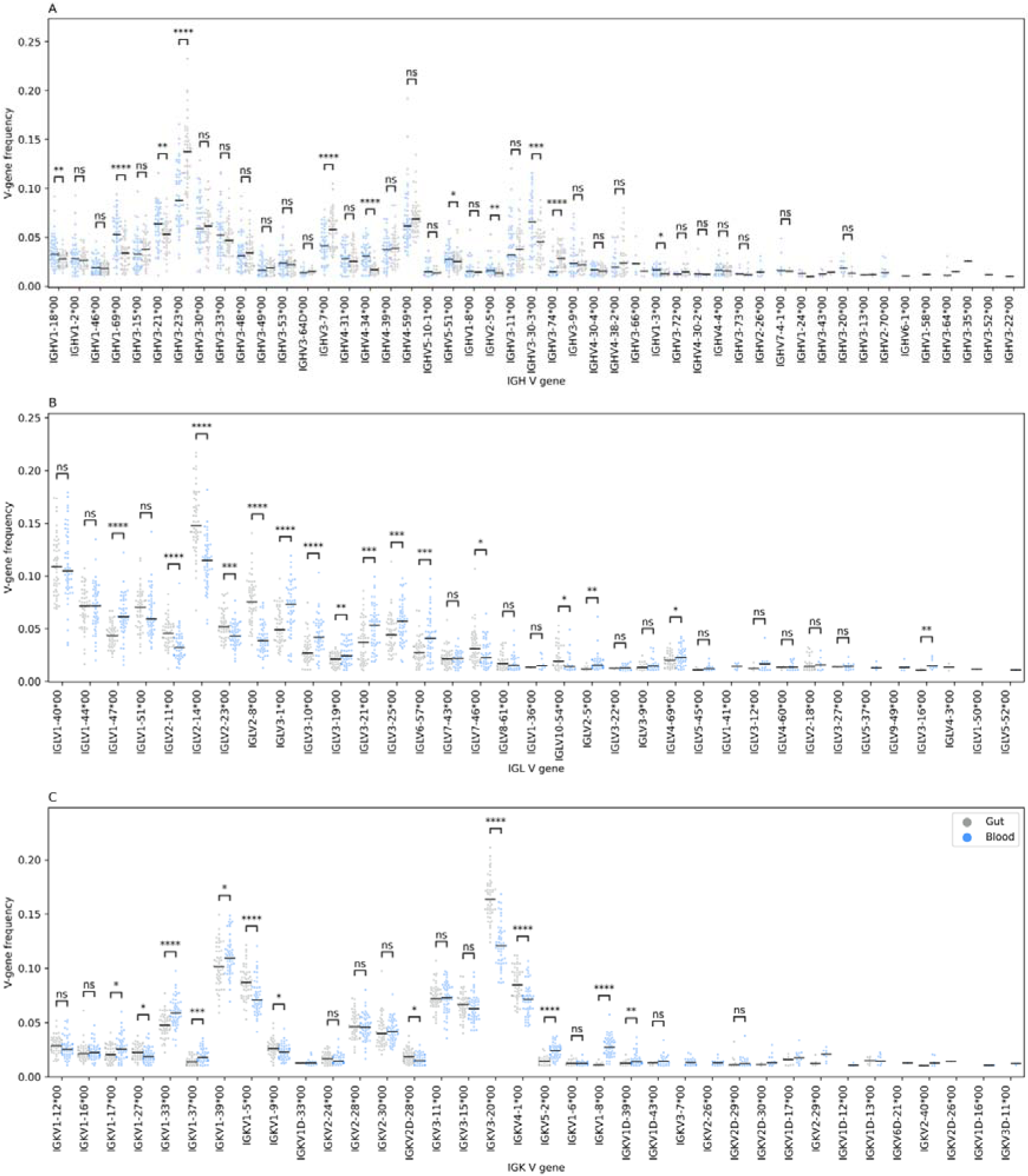
the difference in the V gene usage between the blood and gut B cell receptor chain repertoire. (**A**) shows the difference at the immunoglobin heavy chain (IGH), while (**C**) and (**D**) depict the differences in the V-chain usage for the immunoglobin lambda (IGL) and immunoglobin kappa (IGK) chain, respectively. Across all panels, blackline represents the median, we also used the two-sided Mann-Whitney-Wilcoxon test to compare the frequency of each V-gene segment among the two anatomical compartments. Additionally, we filtered V-genes with a frequency less than 0.01, *i.e.* 1%, from the analysis. Further, we excluded V-gene segments with more than 1% frequency in less than 20 samples from any statistical comparisons.

**Figure S2:**
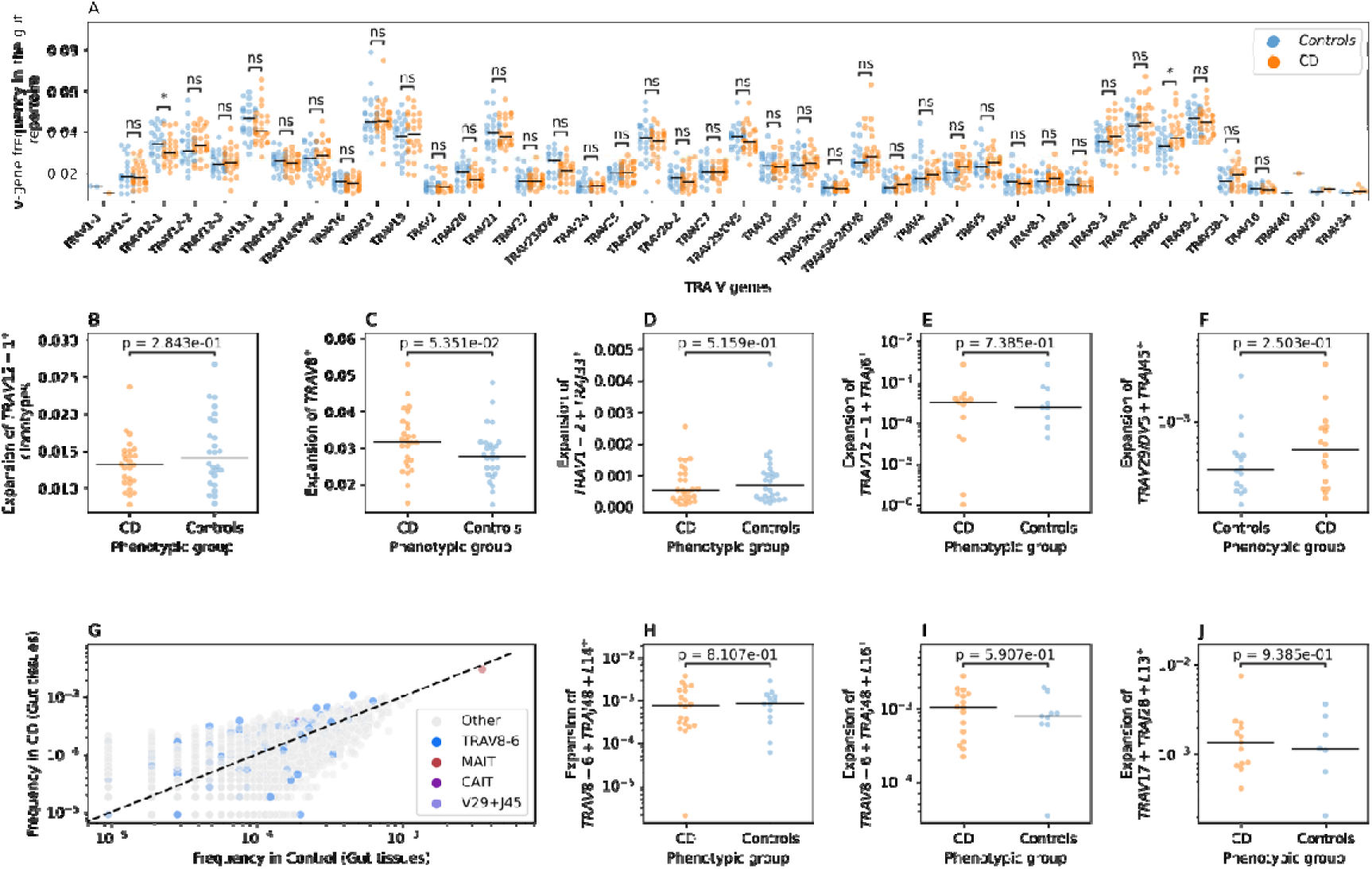
CD-induced changes on the gut TRA repertoire of CD patients and controls. (**A**) depicts a comparison between the frequency of different TRA V-genes in CD patients and controls. (**B**) shows the difference in the expansion of TRAV12-1^+^ clonotypes in CD and Controls, while (**C**) illustrates the same relationship but for TRAV8^+^ clonotypes. (**D**) shows the expansion of the TRAV1-2+TRAJ33+L12 which is used as a proxy for MAIT cells in CD patients and Controls, while (**E**) depicts the same relationship but for TRAV12-1+TRAJ6+L15 group which is a superset of CAIT cells and lastly, (**F**) illustrate the expansion of TRAV29/DV5-TRAJ45+L16 group in the gut of CD patients relative to controls. (**G**) shows the frequency of different VJ-length groups in the TRA gut repertoire of controls and CD patients. (**H**), (**I**) and (**J**) shows the expansion of three VJ-length groups, namely, TRAV8-6+TRAJ48+L14 (**H**), TRAV8-6+TRAJ53+L16 (**I**), TRAV17+TRAJ28+L13 (**J**) in the gut repertoire of CD patients and controls. Across all panels except (**G**), we used the two-sided Mann-Whitney-Wilcoxon test to compare the frequency or the expansion of different V-gene segments or VJ-length groups between CD patients and controls. In (**A**), we filtered V-genes with a frequency less than 0.01, *i.e.* 1%, from the analysis and we excluded V-gene segments with more than 1% frequency in less than 10 samples from any statistical comparisons.

**Figure S3:**
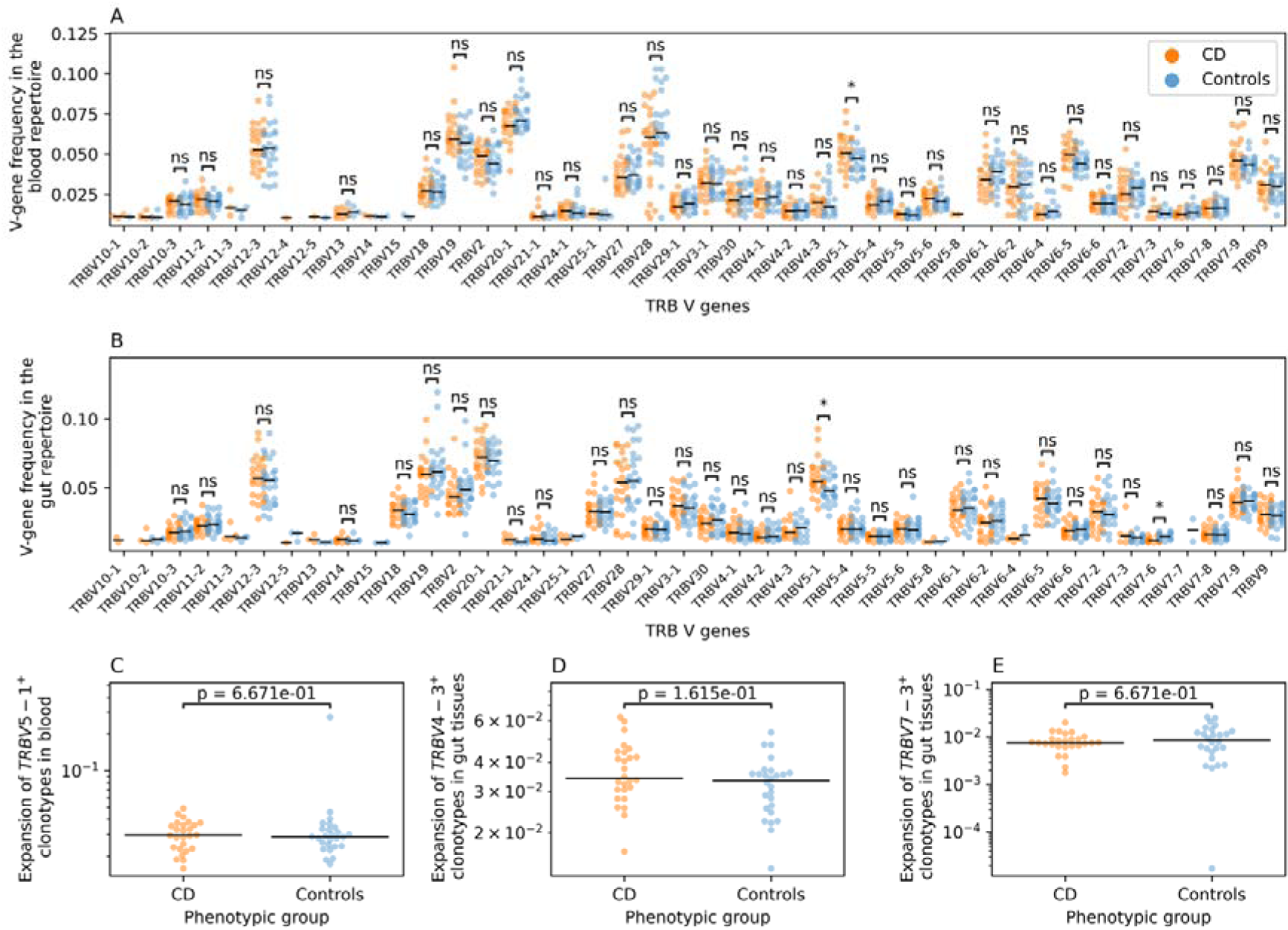
CD-associated changes in the blood and gut TRB repertoire of CD patients and controls. (**A**) and (**B**) depict a comparison between the frequency of different TRB V-genes in the blood (**A**) and gut (**B**) TRB repertoire of CD patients and controls. (**C**) shows the expansion of TRBV5-1^+^ clonotypes in the blood repertoire of CD and controls while (**D**) and (**E**) show the expansion of TRBV4-3^+^ (**D**) and TRBV7-3^+^ (**E**) clonotypes in the gut repertoire of CD patients and controls. Across all panels, we used the two-sided Mann-Whitney-Wilcoxon test to compare the frequency or the expansion of different V-gene segments or VJ-length groups between CD patients and controls. In (**A**) and (**B**), we filtered V-genes with a frequency less than 0.01, *i.e.* 1%, from the analysis and we excluded V-gene segments with more than 1% frequency in less than 10 samples from any statistical comparisons.

**Figure S4:**
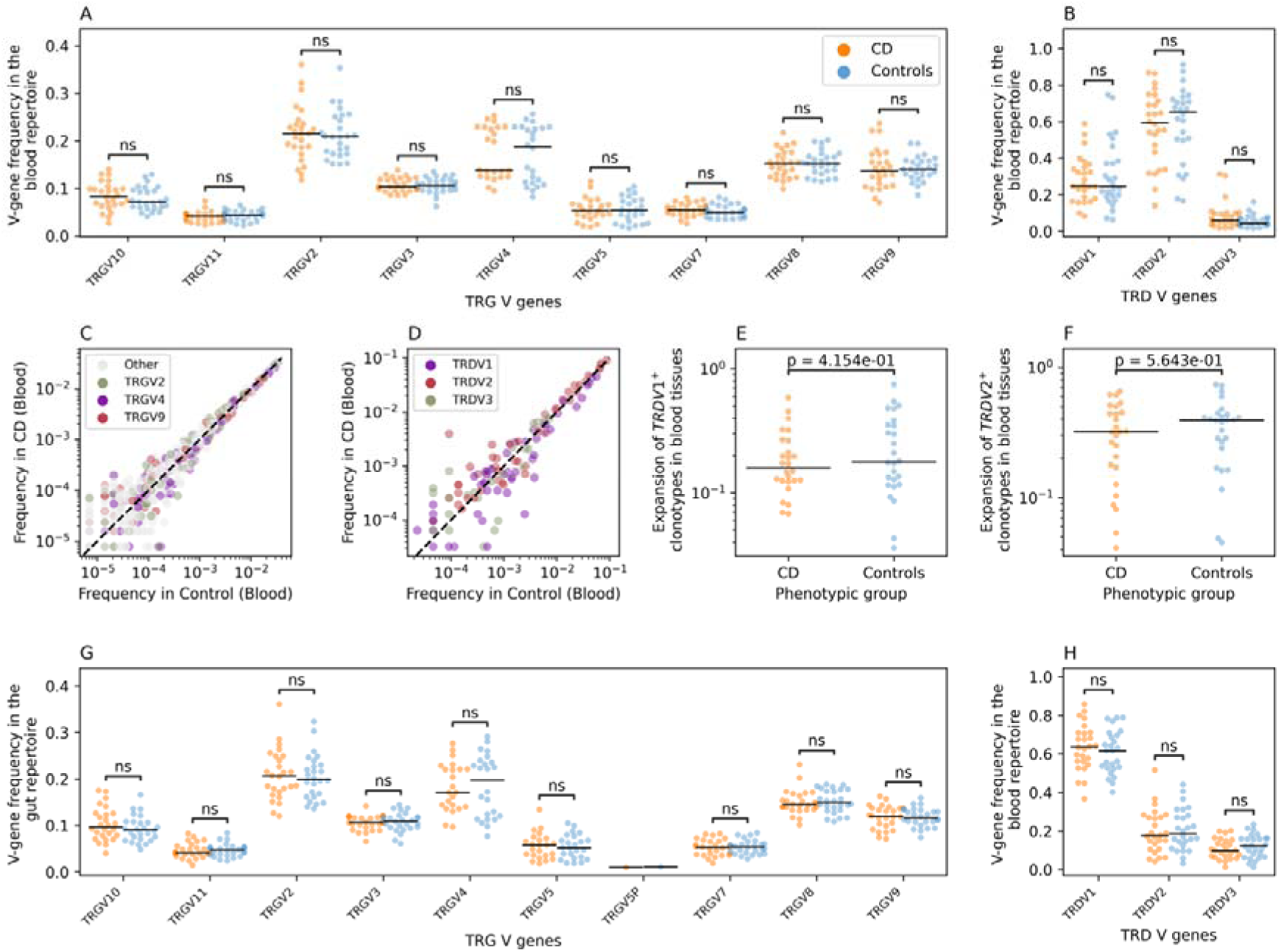
CD-associated changes at the blood and gut TRG and TRD repertoires of CD patients and controls. (**A**) shows a comparison between the frequency of different TRG V-genes in the TRG blood repertoire of CD patients and controls, while (**B**) depicts the same relation but for the different V genes of the TRD blood repertoire. (**C**) and (**D**) show the frequency of different VJ-length groups in the TRG (**A**) and TRD (**B**) blood repertoire of controls and CD patients. (**E**) and (**F**) shows the expansion of TRDV1^+^ (**E**) and TRDV2^+^(**F**) clonotypes in the TRD blood repertoire of CD patients and controls. (**G**) shows a comparison between the frequency of different TRG V-genes in the TRG gut repertoire of CD patients and controls, while (**H**) depicts the same relation but for the different V genes of the TRD gut repertoire. In (**A**), (**B**), (**G**), and (**H**), we filtered V-genes with a frequency less than 0.01, i.e. 1%, from the analysis and we excluded V-gene segments with more than 1% frequency in less than 10 samples from any statistical comparisons. Across all panels, we used the two-sided Mann-Whitney-Wilcoxon test to compare the frequency or the expansion of different V-gene segments or VJ- length groups between CD patients and controls.

**Figure S5:**
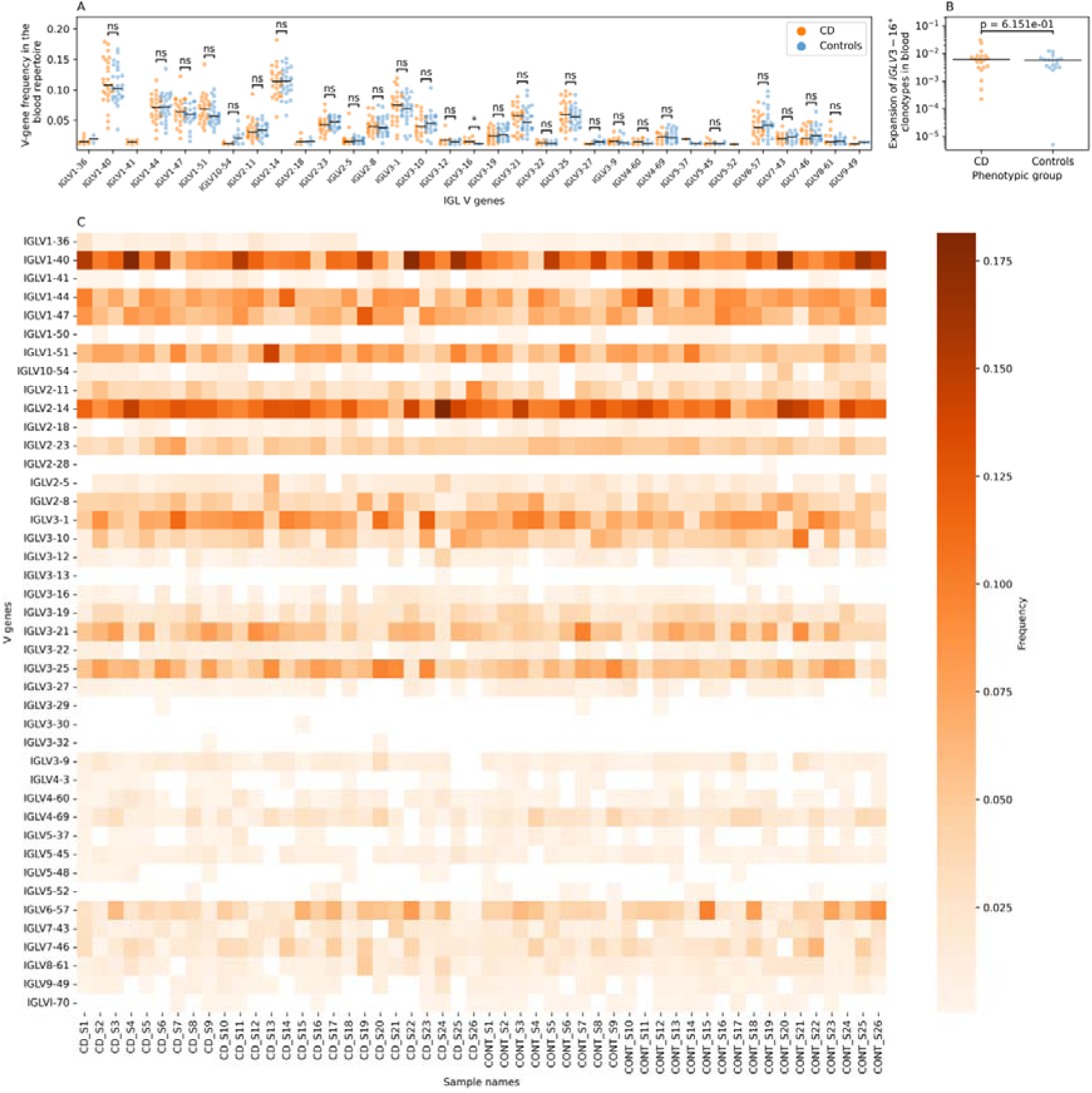
CD-induced changes on the IGL blood repertoire of CD patients and controls. (**A**) shows a comparison between the frequency of different IGL V-genes in the IGL blood repertoire of CD patients and controls. In (**A**) we filtered V-genes with a frequency less than 0.01, *i.e.* 1%, from the analysis and we excluded V-gene segments with more than 1% frequency in less than 10 samples from any statistical comparisons. (**B**) shows the expansion of IGLV3-16^+^ clonotypes in the blood IGL repertoire of CD patients and controls. (**C**) represent a heatmap of the frequency of each V gene in the IGL repertoire of each CD sample (CD_S) and symptomatic control samples (CONT_S) included in the study cohort. In (**A**) and (**B**), we used the two-sided Mann-Whitney-Wilcoxon test to compare the frequency of different V-gene segments (**A**) or the expansion of different VJ-length groups (**B**) between CD patients and controls.

**Figure S6:**
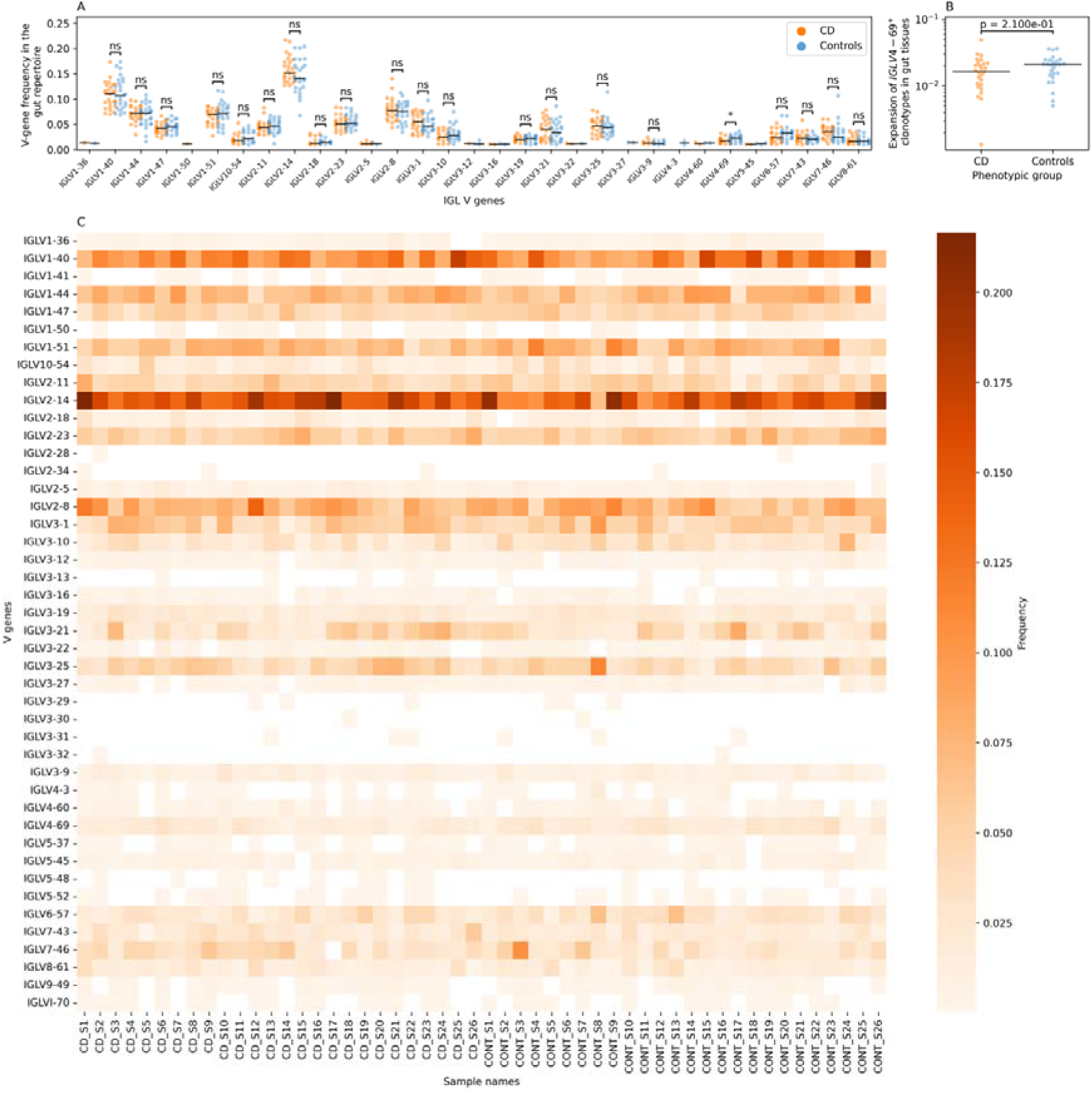
CD-induced changes on the IGL gut repertoire of CD patients and controls. (**A**) shows a comparison between the frequency of different IGL V-genes in the IGL gut repertoire of CD patients and controls. In (**A**) we filtered V-genes with a frequency less than 0.01, *i.e.* 1%, from the analysis and we excluded V-gene segments with more than 1% frequency in less than 10 samples from any statistical comparisons. (**B**) shows the expansion of IGLV4-69^+^ clonotypes in the gut IGL repertoire of CD patients and controls. (**C**) represent a heatmap of the frequency of each V gene in the IGL repertoire of each CD sample (CD_S) and symptomatic control samples (CONT_S) included in the study cohort. In (**A**) and (**B**), we used the two-sided Mann-Whitney-Wilcoxon test to compare the frequency of different V-gene segments (**A**) or the expansion of different VJ-length groups (**B**) between CD patients and controls.

**Figure S7:**
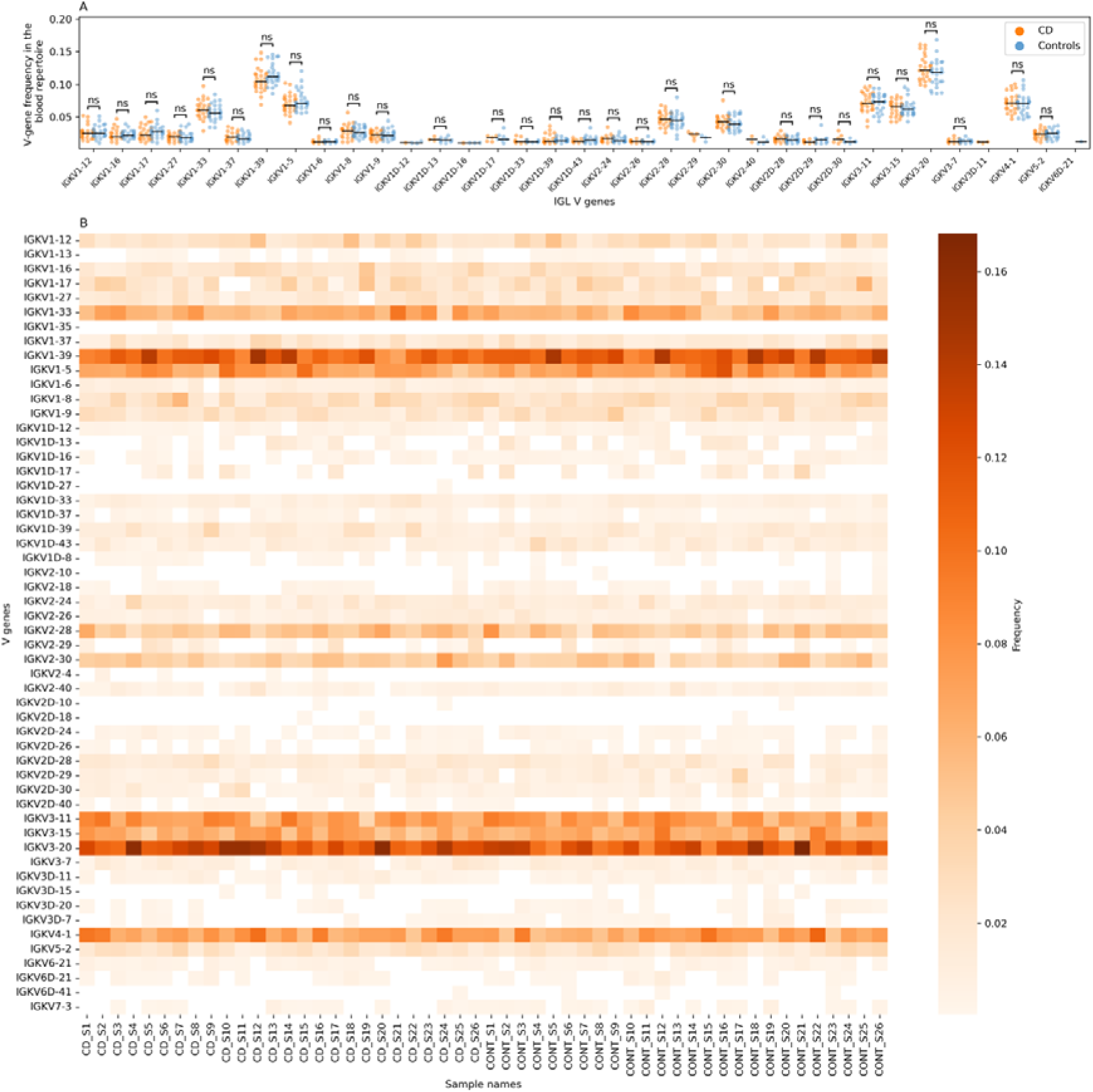
CD-induced changes on the IGK blood repertoire of CD patients and controls. (**A**) shows a comparison between the frequency of different IGK V-genes in the IGK blood repertoire of CD patients and controls. In (**A**) we filtered V-genes with a frequency less than 0.01, *i.e.* 1%, from the analysis and we excluded V-gene segments with more than 1% frequency in less than 10 samples from any statistical comparisons. Also, we used the two-sided Mann-Whitney-Wilcoxon test to compare the frequency of each included V-gene segment between CD patients and controls. (**B**) represents a heatmap of the frequency of each V gene in the blood IGK repertoire of each CD sample (CD_S) and symptomatic control samples (CONT_S) included in the study cohort.

**Figure S8:**
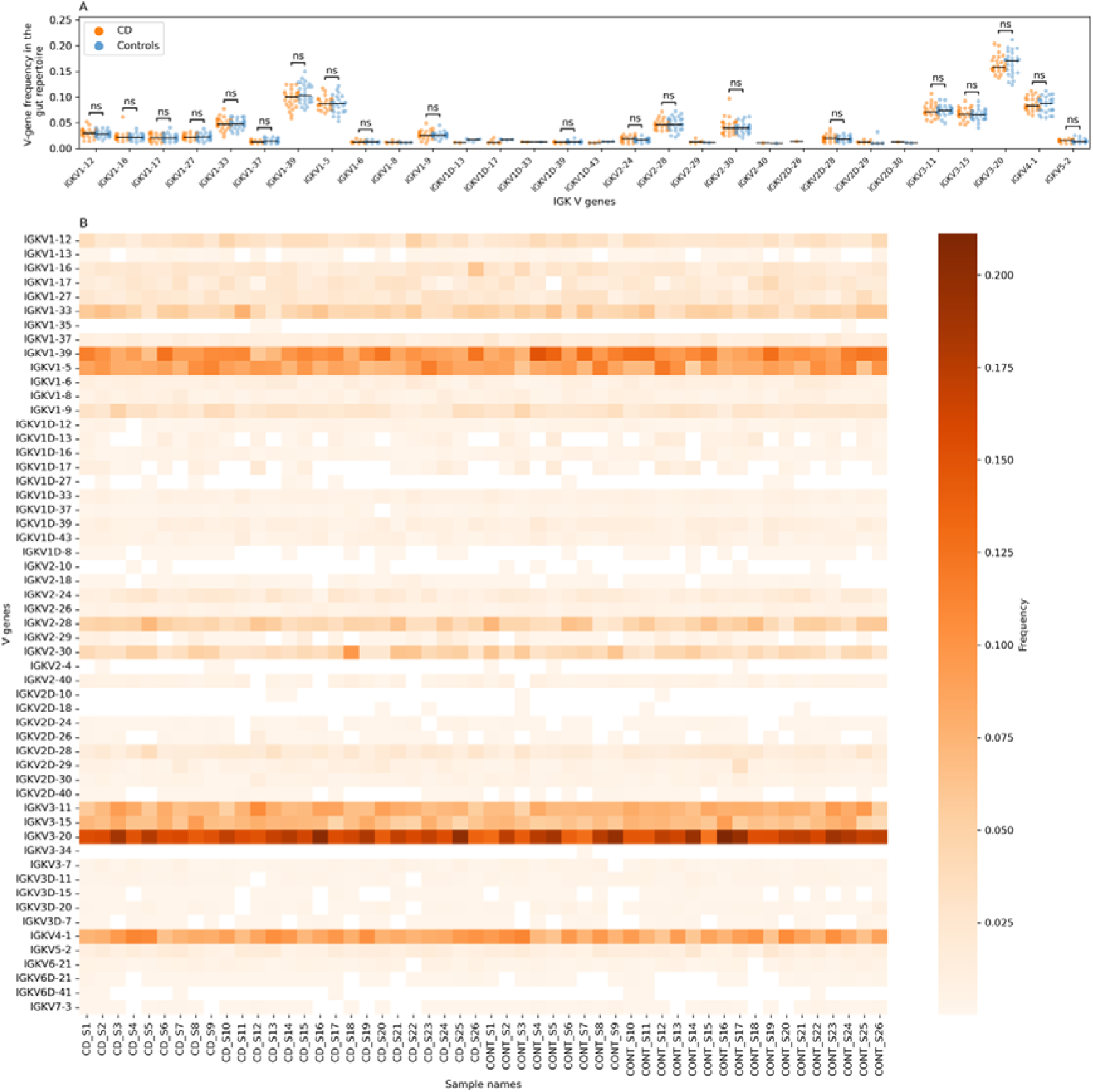
CD-induced changes on the IGK gut repertoire of CD patients and controls. (**A**) shows a comparison between the frequency of different IGK V-genes in the IGK gut repertoire of CD patients and controls. In (**A**) we filtered V-genes with a frequency less than 0.01, *i.e.* 1%, from the analysis and we excluded V-gene segments with more than 1% frequency in less than 10 samples from any statistical comparisons. Also, we used the two-sided Mann-Whitney-Wilcoxon test to compare the frequency of each included V-gene segment between CD patients and controls. (**B**) represents a heatmap of the frequency of each V gene in the gut IGK repertoire of each CD sample (CD_S) and symptomatic control samples (CONT_S) included in the study cohort.

**Figure S9:**
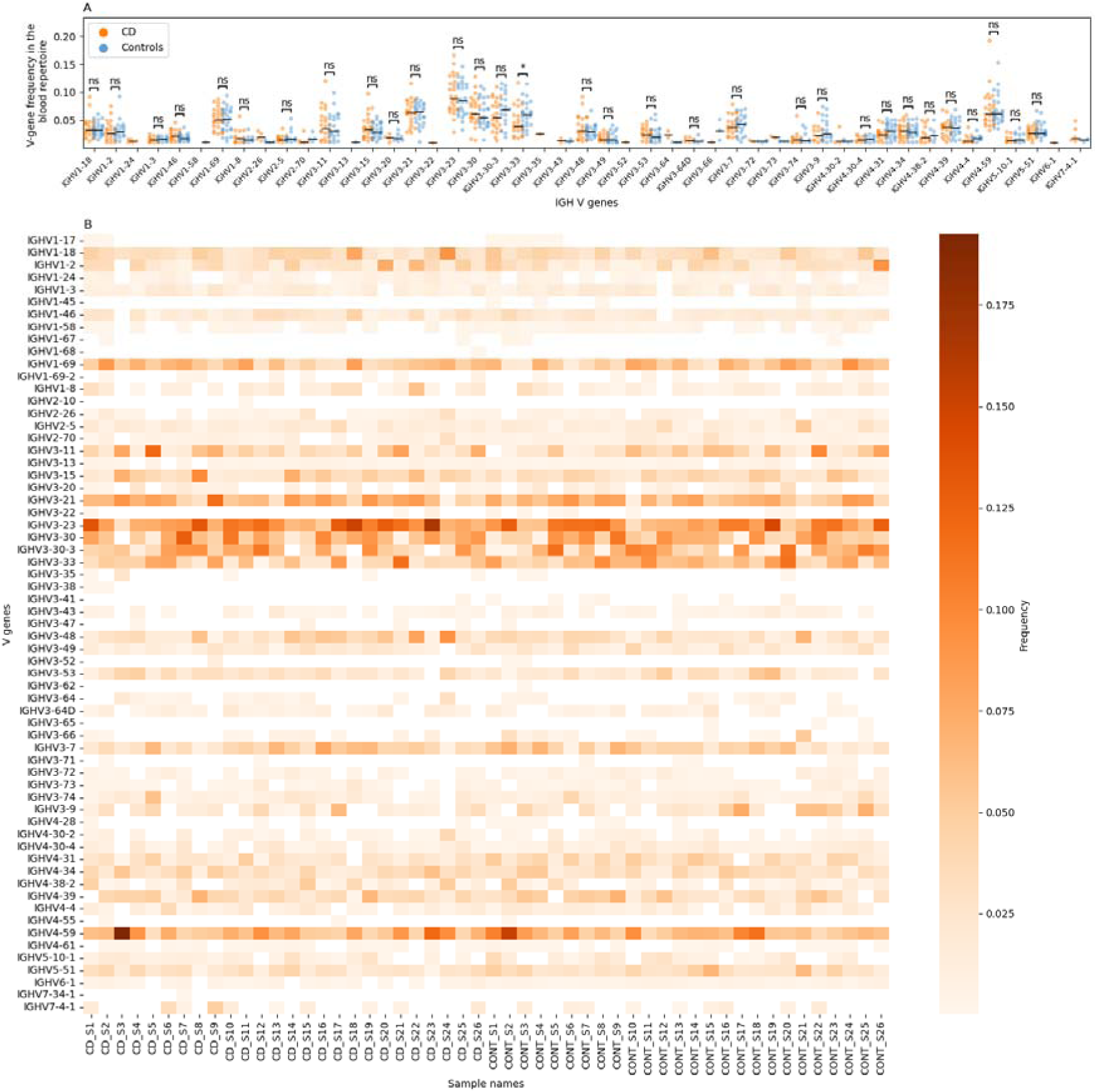
CD-induced changes on the IGH blood repertoire of CD patients and controls. (**A**) shows a comparison between the frequency of different IGH V-genes in the IGH blood repertoire of CD patients and controls. In (**A**) we filtered V-genes with a frequency less than 0.01, *i.e.* 1%, from the analysis and we excluded V-gene segments with more than 1% frequency in less than 10 samples from any statistical comparisons. Also, we used the two-sided Mann-Whitney-Wilcoxon test to compare the frequency of each included V-gene segment between CD patients and controls. (**B**) represents a heatmap of the frequency of each V gene in the blood IGH repertoire of each CD sample (CD_S) and symptomatic control samples (CONT_S) included in the study cohort.

